# Analysis of the SARS-CoV-2 spike protein revealed that blocked receptor-binding domain antigenicity decreases the production of neutralizing antibodies in vivo

**DOI:** 10.1101/2023.02.23.529497

**Authors:** Ya-Fang Mei, Qian Zhang, Maria Hammond, Jie Song, Zongwei Fang, Hongxing Zhao, Marene Landgren

## Abstract

The identification of SARS-CoV-2 spike protein distribution and function in target cells has raised concerns about its possible impacts on vaccine efficacy and pathogenic effect in host cells. Thus, a better understanding of such consequences is necessary. In this study, we studied the biological characteristics of six variants of SARS-CoV-2 in A549 and HEK293 cells using four different technologies. The results showed that compared to the other fragments, the full-length spike protein exhibited the highest expression on the cell surface and was detectable in the cell supernatant, cytoplasm, and nucleus. Except for the cell surface, the S1 subunit generally expressed higher than the full-length spike protein. RBD and S2 subunits were expressed in the cytoskeleton. The SS-RBD peptide, which consists of a 19-amino acid signal peptide sequence (SS)-linked RBD, exhibited the highest expression in the cell supernatant among all other studied peptides. The SS positively enhanced the expression, migration, and secretion of SS-RBD from the cytoskeleton to the supernatant. Importantly, the FACS assay results showed that neutralizing antibodies (NAbs) could recognize SS-RBD but not RBD in the transfected cells, suggesting that RBD was tightly bound by ACE2 in HEK293 cells. In contrast, the antigenicity of the RBD in the spike protein was revealed and efficiently monitored only by 6-His-tag mAbs. Thus, our findings demonstrated that ACE2 blocks crucial immunogenic epitopes of the RBD, and the full-length spike protein mainly induces non-neutralizing antibodies in vivo. Therefore, we suggest that reducing ACE2 binding affinity and exposing the immunogenicity of the RBD on the spike protein is imperative for improving vaccine efficacy and generating new SARS-CoV-2 mRNA vaccines.

## Introduction

Severe acute respiratory syndrome coronavirus 2 (SARS-CoV-2) negatively influences global public health, economic development, and social activities. Since the pandemic outbreak, unprecedented efforts have been made to develop SARS-CoV-2 vaccines that can be used against the pandemic worldwide. Usually, the spike protein is one of the main structural proteins that plays a crucial role in eliciting immune responses during disease progression. Therefore, it is considered a major antigenic protein that can induce neutralizing antibodies (NAbs) ^1^. Moreover, the highly glycosylated spike protein ^2^ is also fundamental for viral pathogenesis, transmission, and virulence because it binds to the human host cell membrane by interacting with the host angiotensin-converting enzyme 2 (hACE2) receptor ^3–5^. Wang QH *et al*. reported the atomic structure of the receptor-binding domain (RBD) interface and ACE2 receptors^6^. Simultaneously, potent NAbs are elicited and competitively bind to the ACE2 receptor to block the interaction between the RBD and ACE2 receptor. Some NAbs outside the receptor binding motif (RBM) have also been identified ^7^.

To date, several COVID-19 vaccines have been generated based on the genomic sequence of the spike protein (Wu *et al*., 2020). Based on the severity of SARS-CoV-2, vaccine prevention has positively impacted society ^8^. Population-level immunity against SARS-CoV-2 has substantially reduced the number of COVID-19 hospitalizations and mortality rates ^9 10^. However, new Omicron variants are emerging all over the world. Moreover, adverse postvaccination effects, such as SARS-CoV-2 reinfection, myocarditis, pericarditis, and multiple necrotic encephalitis, have been reported ^11,12^. Although several spike protein structures and their receptor binding sites are well-known ^13^, more information is needed concerning the biology of spike proteins in transfected cells. Highly contagious SARS-CoV-2 viruses with different variants of the spike protein have started emerging since the beginning of the COVID-19 pandemic. Different spike protein variants may cause biological, antigenic, and pathogenic changes ^14,15^. Our group and other researchers have previously reported the subcellular distributions of the SARS-CoV-2 spike protein ^16,17^. It is essential to thoroughly investigate the location and expression of the spike protein and its fragments in vitro to gain a deeper understanding of the interaction between spike proteins and host cells. Moreover, this research provides insights into tracking spike proteins in cellular compartments. In this study, we used the full-length spike protein’s biological distribution and immunological function to identify highly expressed fragments and evaluated potential advantages and disadvantages in the context of vaccine development.

## Materials and Methods

### Construction of recombinant spike protein and variants

The SARS-CoV-2 spike protein sequence was obtained from the GenBank database under accession number MN908947. Spike protein and fragment cDNA were synthesized with codon optimization and cloned into a mammalian expression vector pUC57 with a C–terminal 6xHis tag. In addition, the S1 subunit from the original cDNA without codon modification was generated and studied in parallel as S1-ori.

### Cell lines and antibodies

HEK293 cells were purchased from Microbix Biosystems Inc., Toronto, Canada, and A549 cells were provided by Walter Nelson-Rees (UC Berkeley, USA). The cell culture and passage of HEK293 and A549 cells were carried out as previously described (Mei *et al*., 2016). A rabbit anti-SARS-CoV-2 neutralizing antibody (4G6) was purchased from Genscript (Cat. No. A02053-100). Rabbit monoclonal antibodies against 6-His-tags (12698S) were purchased from Cell Signaling Technology. The rabbit monoclonal anti-histone H2A. X antibody (ab124781), calreticulin ER marker (ab92516), cytoskeleton marker (Cytoskeleton 18, ab133263), β-tubulin antibody (ab6046), and goat anti-rabbit IgG H & L HRP antibody (ab207718) were purchased from Abcam. A FITC-conjugated polyclonal swine anti-rabbit immunoglobulin antibody purchased from DAKO and a nonconjugated normal rabbit IgG control antibody purchased from Santa Cruz Biotechnology (sc-3888) were also used.

### FACS assay

For kinetics experiments, 3×10^5^ cells were seeded in a 12-well plate and grown overnight. The next day, cells were transfected with the spike protein plasmids. According to the manufacturer’s protocol, the manufacturer mixed 1,6 or 1,1 μg/well of DNA from each plasmid with 4 μl of Lipofectamine 2000 and cultivated for 24, 48, or 72 hrs. Then, the cells were detached in PBS-EDTA buffer and centrifuged at 800 rpm for 5 min; the cells were washed once with 1% bovine serum albumin (BSA) PBS (BP). The cells were resuspended in 100 μl of 4% paraformaldehyde for 20 min at RT and washed once in BP at 4°C. The cells were then incubated with rabbit anti-spike protein polyclonal antibody (NAb) (diluted 1:1000 in BP) for one hr at 4°C with gentle shaking. After one hr of incubation, the cells were washed once with the same buffer and incubated with FITC-conjugated polyclonal swine anti-rabbit immunoglobulin antibody from DAKO (dilution 1:40 in BP) for 1 hr at 4°C on ice with shaking. Afterward, the cells were washed with BP and analyzed by flow cytometry using a Bio-Rad ZE5 instrument (USA). The results were analyzed using Hacking Flowjo VX software (EXCITE, EXPERT cytometry, Design: BKSURU.com. USA). In addition, considering that detection of the RBD might block as a result of ACE2 interaction, a rabbit monoclonal anti-6-His tag antibody was also used to detect RBD expression in the FACS assay. Cells transfected with an empty vector were used as the negative control.

### Cell lysate, pellet, and supernatant preparation

The culture medium was collected from the transfected cells and centrifuged at 1500 rpm for 5 min to pellet the cells. The supernatant was collected. The cell pellet was washed once with cold PBS. The cell lysate and pellet were prepared according to the protocol of Bio-Rad (www.biorad.com/webroot/web/pdf/lsr/literature/Bulletin_6376.pdf). All the samples were stored at −80°C. The protein concentration of the cell lysate was measured by BCA assay (Pierce BCA Protein Assay Kit, Thermo Scientific, USA). The cell pellet was resuspended in 40 μl ddH_2_O and mixed carefully. The cell supernatant was harvested at 24, 48, and 72 hrs posttransfection (p.t.). The harvested cell supernatant was immediately frozen at −80°C. Proximity extension assays (PEA), PCR experiments, and immunoblot assays were used to assess the secreted levels of the full-length spike protein and fragments in the cell supernatant.

### SDS–PAGE and immunoblot assay

SDS–PAGE and Western blotting were used to detect the SARS-CoV-2 spike protein and fragments in the cell lysates, pellets, and supernatants; thirty microliters of supernatant, 10 μg of cell lysate, and 5 μl of resuspended cell pellet solution were loaded into each well. Proteins in the samples were separated in NuPAGE SDS–PAGE gels and transferred to nitrocellulose membranes according to the protocol from the NuPAGE Technical Guide with minor modifications (www.biorad.com/webroot/web/pdf/lsr/literature/Bulletin_6376.pdf). After the membrane was blocked with 5% BSA (Bovine Serum Albumin Fraction V, Roche Diagnostics, Germany) in TBST buffer (1X Tris-Buffered saline, 0.1% Tween-20 detergent, 20 mM Tris-HCl, 150 mM NaCl, 0.1% Tween 20, pH=7.4), the membranes were incubated with anti-His-tag primary antibodies (1:2000) overnight at 4°C, followed by incubation with horseradish peroxidase (HRP)-conjugated secondary antibodies (Goat pAb to Rb IgG, Abcam, 1:10000). Blotted nitrocellulose membranes were visualized with ECL Reagent (Thermo Fisher, USA) and CLXposure Film (Thermo Fisher, USA) and the exposure time ranged from 1 sec to 30 min.

### Immunoblotting for subcellular fractionated proteins

A total of 1×10^6^ cells in a 6-well plate were transfected with five various spike protein fragments. At 48 hr posttransfection (p.t.), cells were fractionated using a subcellular protein fractionation kit (Thermo Scientific cat. no. 78840). Eluted proteins from subcellular fractions were quantified by a Pierce ™ BCA protein assay kit (Cat.no: 23227. Thermo Fisher Scientific, USA. The normalized concentration of each extract was 10 μg, but the concentration of the cytoskeleton fraction was 5 μg due to the limited number of proteins obtained in this fraction. The samples were analyzed by Western blotting using specific antibodies against proteins from various subcellular fractions. The primary antibodies included calreticulin (Calreticulin ER marker: ab92516) for detecting membrane (M) and endoplasmic reticulum (SN) fractions, beta-tubulin for detecting the cytoplasmic fraction (CP) (beta-tubulin, ab6046), cytoskeleton marker (Cytoskeleton 18, ab133263) for detecting the soluble nuclear (SN) and cytoskeleton (CS) fractions, and rabbit monoclonal anti-histone H2A. X Ab (ab124781) for detecting the chromatin-bound nuclear (CBN) fraction. The spike protein and fragments were detected by rabbit monoclonal antibodies (mAb) against the 6-His tag (12698S) from Cell Signaling Technology. The nitrocellulose (NC) membrane was immunoblotted with specific primary antibodies, followed by incubation with HRP-conjugated secondary antibodies (Abcam). The dilutions for primary antibodies were 1:1000, and the secondary antibody was 1:10000. Protein bands were detected by Pierce ECL Western Blotting Substrate (Thermo Fisher Scientific, Cat no, 32106).

### Proximity extension assays (PEA)

The full-length spike protein and fragments produced from HEK293 cells were further evaluated by highly sensitive proximity extension assays (PEA)^18,19^. Thirty micrograms of rabbit anti-SARS-CoV-2 (2019-nCoV) spike-RBD polyclonal antibody (catalog no: 40592-T62. Sinobiological, US) was activated with dibenzylcyclooctyne-NHS ester (DBCO-NHS ester, Sigma Aldrich) at room temperature (RT) for 30 min. The unreacted DBCO-NHS ester was removed using a 7K MWCO Zeba desalting spin column (Thermo Scientific). Activated proteins were divided into two aliquots, and each aliquot was incubated at 4°C overnight with one pair of azide-modified oligonucleotides ^20^. PEA was performed as previously described. Briefly, a pair of 1.33 nM oligonucleotide-conjugated antibodies were incubated with 1 μl or dilutions thereof of activated proteins in medium collected at 24, 48, and 72 hr p.t. at 37°C for 1 hr or 4°C overnight. The pair of oligonucleotides is brought in proximity via the interaction of the antibody with the S1 protein. After incubation, 96 μl of extension solution containing 1X Hypernova buffer (BLIRT S.A.), 1.5 mM MgCl_2_, 0.2 mM of each dNTP, 1 μM of the universal forward and reverse primers, 2 U/ml DNA polymerase (Invitrogen) and 5 U/ml Hypernova DNA polymerase was added. The extension/pre-PCRs were run at 50°C for 20 min, followed by a 5-min heat-activation step at 95°C and 17 cycles of pre-PCR of 95°C, 30 sec, 54°C, 1 min, and 60°C, 1 min. For the subsequent qPCR detection, 2.5 μl of the extension/pre-PCR products were transferred to a 96 or 384-well plate and combined with 7.5 μl qPCR mix containing 1X PCR buffer (Invitrogen), 2.5 mM MgCl_2_, 0.2 mM of each dNTP, 1.6 μM ROX reference Dye (Invitrogen), 0.3 μM molecular beacon (Biomers), 0.9 μM of each primer, 0.1 U Uracil N-glycosylase and 0.5 U recombinant Taq polymerase (Invitrogen). Quantitative real-time PCR was run with initial incubation at 25°C for 30 min, then denaturation at 95°C for 5 min, followed by 30 cycles of 15-sec denaturation at 95°C and 1 min annealing/extension at 60°C. All PCR primers and the molecular signal used in this study have been previously published^19^. The RBS, SS-RBD, and S1 fragments and the full-length spike protein, but not the S2 fragment, were detected in these assays.

### Immunofluorescence and confocal microscope

A549 and HEK293 cells were plated on 12-mm coverslips overnight in a 24-well plate at a density of 80,000 cells/well. Cells were transfected with Lipofectamine 2000 and five DNA plasmids or empty vector as a control for 24 hr at 37°C. The cells were washed in PBS and fixed using 4% paraformaldehyde (PFA) in PBS for 20 min at RT. The cells were extensively washed with PBS and permeabilized in 0.5% Triton X-100 for 10 min. The cells were blocked in 5% BSA and then labeled with anti-6-histag mAb diluted in 1% BSA-PBS overnight at 4°C. The cells were washed and labeled with 1:40 diluted swine anti-rabbit immunoglobulin FITC-conjugated secondary polyclonal antibody in 1% BSA-PBS at RT for 1 hr. After that, they were stained with DAPI for 1 min at RT and extensively washed. Coverslips were mounted using Dako Fluorescence Mounting Medium (Agilent) and imaged using a Nikon confocal microscope (Eclipse C1 Plus). All scoring was performed under blinded conditions. Photomicrographs were obtained by an LSM 710 confocal microscope with a 63x objective lens (numerical aperture 1.4) (Carl Zeiss) and visualized by Zen 2011 software. The images were acquired under oil immersion at RT.

### Statistical analysis

Statistical analyses (t-tests and one-way ANOVA) were performed with GraphPad Prism Software version 7.0 (GraphPad Software, San Diego, CA, USA).

## Results

### The spike protein and fragments were significantly expressed in lung and kidney cells

Based on the first reported genome sequence of the SARS-CoV-2 virus (GenBank: MN908947.3), we constructed full-length spike proteins and fragments and added a 6-histidine-target in the C-terminus. A schematic of the constructs can be seen in Fig. 1A, B. The constructs with codon optimization for human mammalian cells were synthesized, and an original S1 fragment (S1-ori) was also synthesized as a control. Then, they were cloned into the mammalian expression vector pUC57 vector (Genscript). The expression cassette consisted of a CMV promoter, spike protein fragments of various sizes, a C-terminal 6xHis-tag, a stop codon, and a SV40 poly-A sequence. A 19 amino acid sequence, called the signal peptide sequence (SS), existed in all but the RBD and S2 fragments (Fig. 1A). The spike glycoprotein ectodomain retained its original amino acid sequence with no mutation in the furin-like cleavage site. Computer programs predicted secondary structure, hydrophilic and hydrophobic domains, and antigenic epitopes on the full-length spike protein (Fig. 1B). The analysis revealed that the full-length spike protein displayed many antigenic epitopes and hydrophilic domains but limited hydrophobic domains.

**Figure 1.**
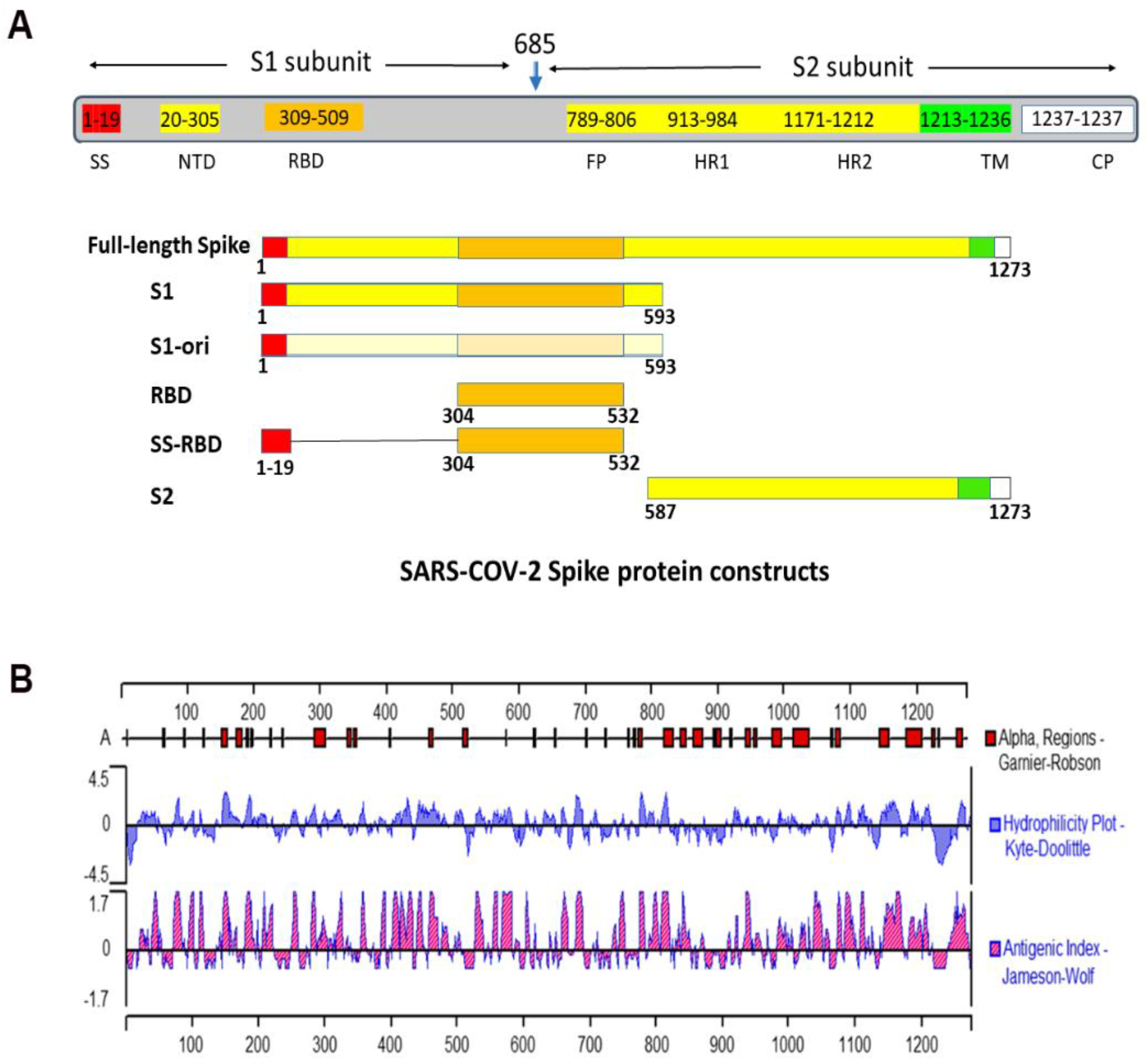
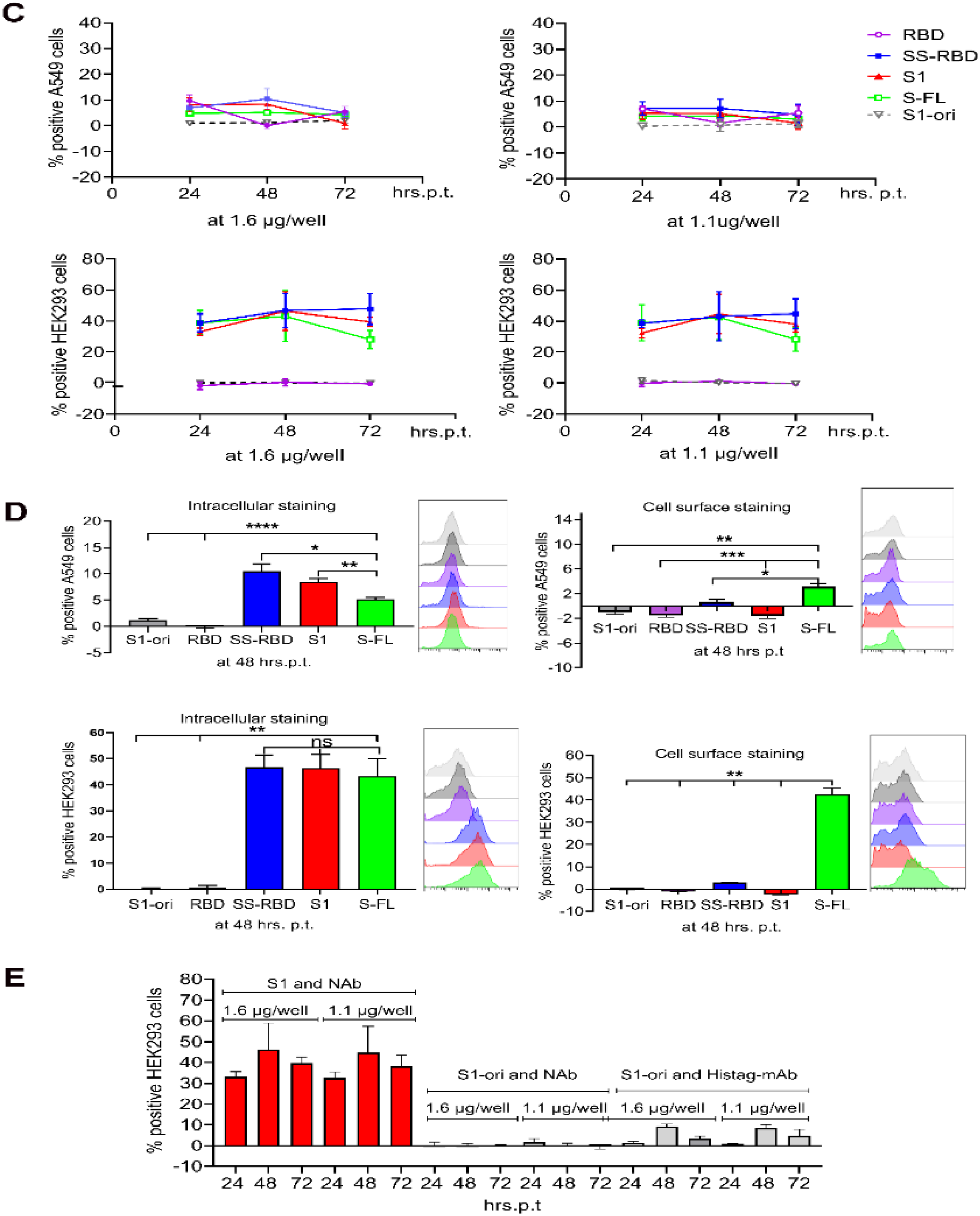
Schematic diagram of the SARS-CoV-2 spike protein and its subdomains expressed in A549 and HEK293 cells. **Fig. 1A:** The full-length spike protein is a heavily glycosylated trimer, and each protomer comprises 1273 amino acids (aa). The demarcation at position 685 represents subunits 1 (S1) and 2 (S2). Subunit S1 is composed of 685 aa and organized into two domains: an N-terminal domain (NTD) and a receptor-binding domain (RBD). The transmembrane S2 subunit is composed of 588 aa (residues 686-1273). It contains an N-terminal hydrophobic fusion peptide (FP), two heptad repeats (HR1 and HR2), a transmembrane domain (TM), and a cytoplasmic tail (CP) ^13^. This article uses spike fragment locations and precise sizes: full-length spike protein (1-1273 aa), S1 (1-593 aa), RBD (304-532 aa), SS-RBD (N-terminal 19 aa plus RBD), and S2 domain (587-1273 aa). The receptor binding motif (RBM) encompasses residues 437 to 508. All protein constructs contained a 6X His-tag at the C-terminus. Signal peptide sequence (SS). A glycan site, “NLT,” is marked with a blue highlight. **Fig. 1B:** Computer programs predict the full-length spike’s secondary structure, hydrophilic and hydrophobic, and antigenic epitopes. **Fig. 1C:** Spike and fragment kinetic expression was detected with intracellular staining by a neutralization antibody (NAb) against the S1 subunit and flow cytometry-based quantification. The empty pUC57 vector was used as a background control. Normal rabbit serum was used as an antibody control. **Fig. 1D:** The expression levels of the full-length spike protein and fragments were evaluated by intracellular staining and cell surface staining with a NAb against the S1 subunit. **Fig. 1E:** Kinetic expression of S1-ori was detected by mAb against 6-histag in FACS analysis and intracellular staining. S1-ori is the original sequence, and the S1 subunit is a humanized genomic sequence. The results indicated that the S1 subunit shows much higher expression than S1-ori. All experiments were performed three times with duplicate samples in each experiment. Error bars represent the mean ± SD. **P < 0.01; ***P < 0.001.

A polyclonal NAb was used in FACS analysis. Two doses at 1.1 μg or 1.6 μg per plasmid were used for parallel expression efficacy studies. The SS-RBD and S1 fragments and the full-length spike protein showed similar efficiency, with an expression rate of approximately 40% in HEK293 cells at 24 hrs p.t. (Fig. 1 C). Then, the expression rate increased from 45 to 55% at 48 hrs. p.t., and SS-RBD showed the highest expression on day 3. S1 and the full-length spike protein exhibited slightly reduced expression rates to approximately 40% positive cells. Next, we investigated the surface expression rate in transfected cells. The full-length spike protein had the highest expression rate on the cell surface of HEK293 and A549 cells among all samples studied. HEK293 cells were 40% positive at 48 hrs p.t. (Fig. 1D). In general, the expression of spikes or fragments was lower in A549 cells than in HEK293 cells. Moreover, the two doses of plasmids provided comparable values in the FACS assay. As a result, we chose to use 1.6 μg per well in the following experiments.

S1-ori showed an expression rate of approximately 1%, according to the FACS analysis. To verify the data, we repeated the experiments using an anti-6-Histag mAb instead of a NAb and compared S1-ori and S1 with genomic optimization for human cells. We found that S1-ori was expressed at a minimal level using either mAb or NAb in the FACS assay. Therefore, we assumed that S1-ori was unsuitable for vaccine purposes (Fig. 1E). All spike proteins and variants used in the following experiments were necessarily optimized for gene expression in human cells.

### Anti-6-his tag mAb effectively detected RBD expression

The FACS assay results showed that the RBD, which is critically involved in ACE-2 binding and neutralizing antigens, was undetectable in transfected cells. We considered that the two functional domains could overlap together, resulting in the blocked antigens escaping the detection of NAb. We switched the NAb to the 6-his tag mAb and repeated intracellular staining for RBD expression. As expected, the 6-histag mAb effectively detected RBD expression. We noted that the SS-RBD fragment could be observed in the same assay. We speculated that SS could impact the confirmation of upstream antigenic epitopes of RBD, resulting in the α1 epitope being revealed and the NAb binding it. This study used the first published spike protein sequence to construct the available DNA plasmids. To elucidate the interaction between RBD and ACE-2, we adapted the three-dimensional structure of RBD-ACE2 reported by Wang *et al*. ^6^. Overall, we found that the ACE-2 receptor blocked the antigenicity of the RBD in host cells (Fig. 2D-E.). Moreover, we also remarked on the critical amino acids in Fig. 2F.

**Figure 2.**
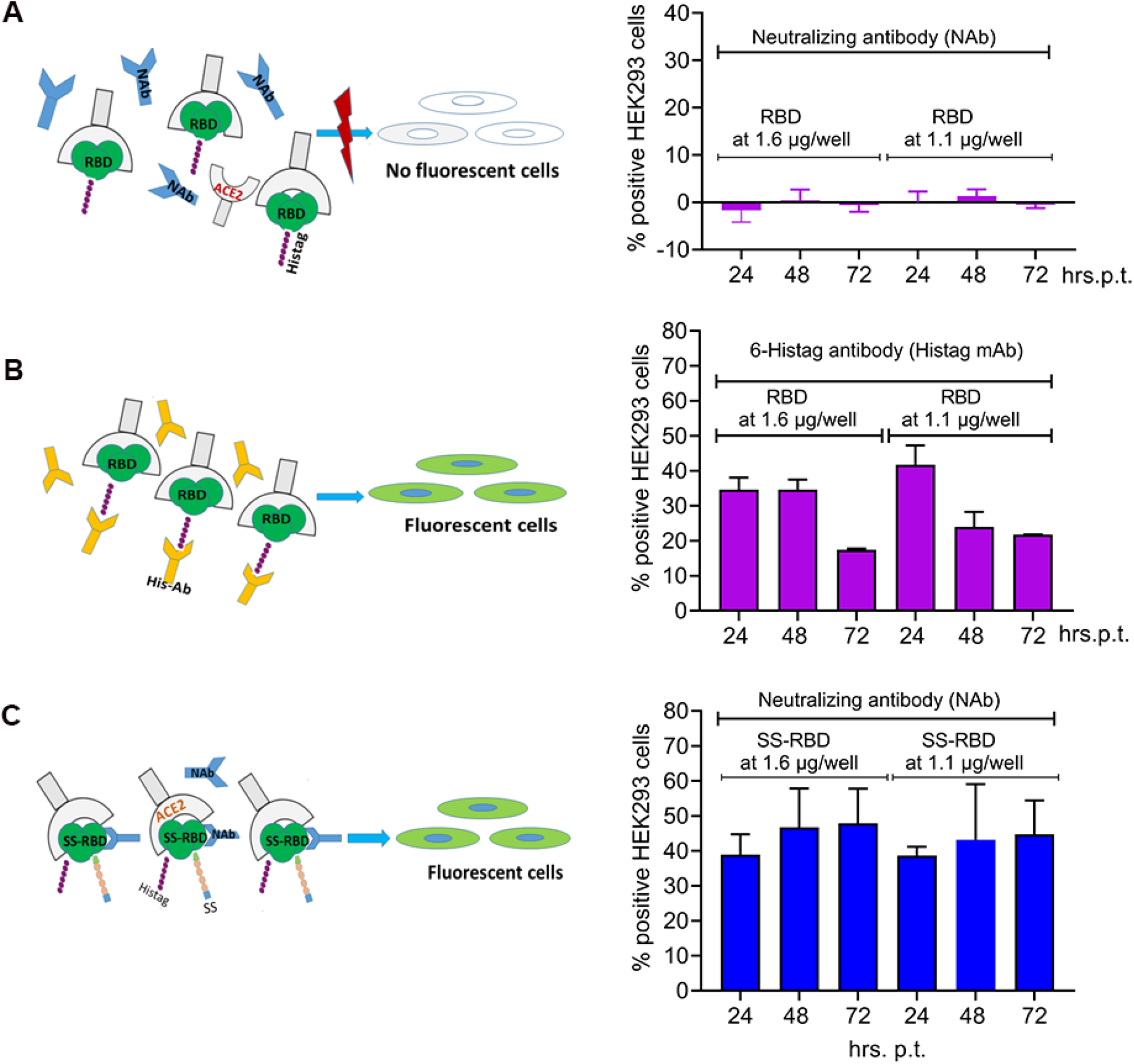
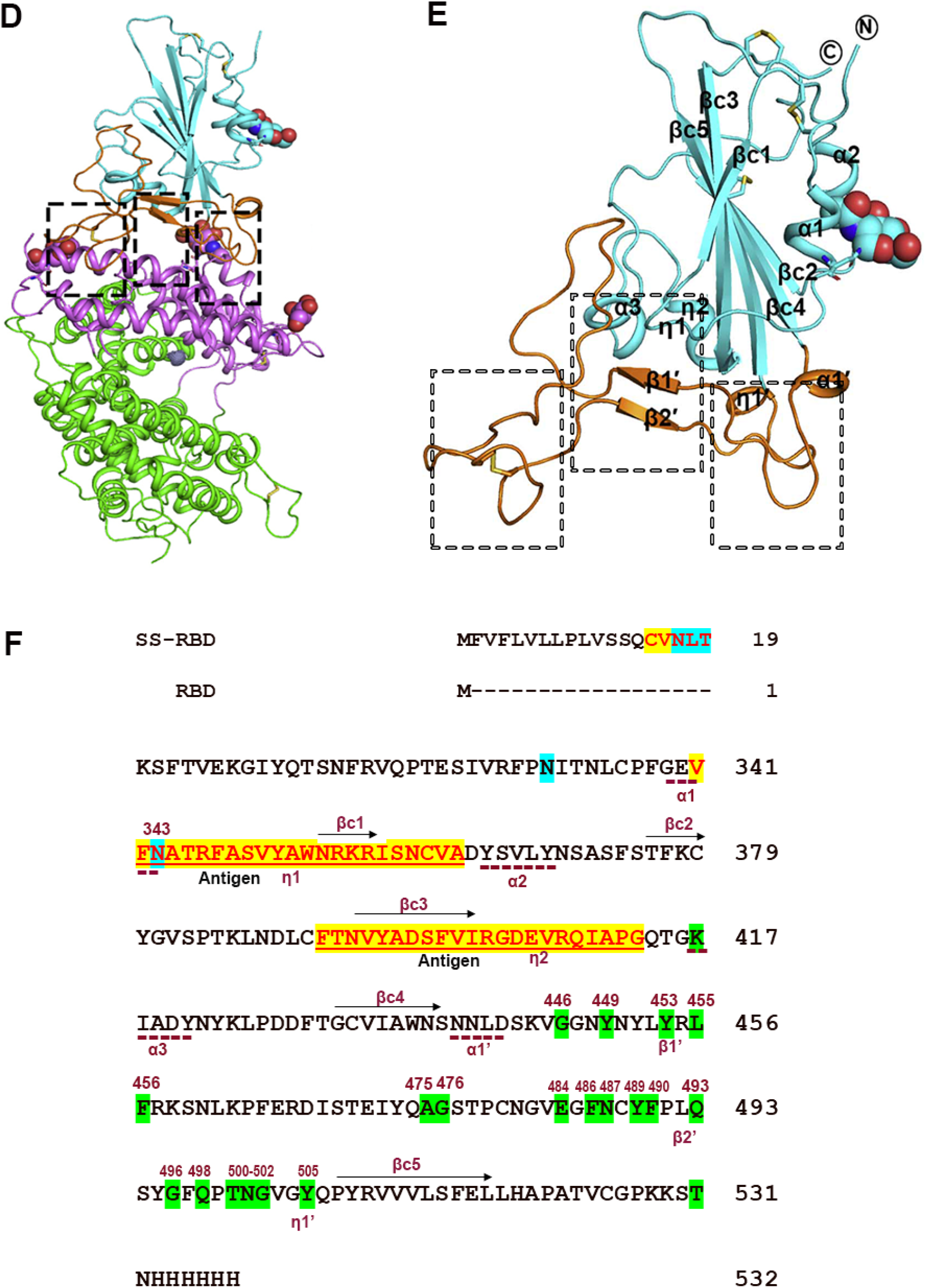
Illustrations of the interaction between RBD/SS-RBD and ACE2 receptor and the mechanism by which RBD is undetectable with a NAb but detectable with an anti-6-Histag mAb. **Fig. 2A:** Kinetics in RBD expression were detected by intracellular staining with a NAb against the S1 subunit in HEK293 cells. RBD plasmids administered at 1,6 and 1,1 μg/well were studied in parallel. The empty pUC57 vector was used as a negative control. **Fig. 2B:** Kinetics of RBD expression were detected by FACS analysis with intracellular staining and anti-6 His-tag mAb. **Fig. 2C:** Kinetics of SS-RBD expression were detected with intracellular staining by NAb against the S1 subunit. **Fig. 2D:** The complex structure of SARS-CoV-2 RBD bound to hACE2. The core and external subdomains in the SARS-CoV-2 RBD are colored cyan and orange, respectively. Human ACE2 subdomains I and II are colored violet and green, respectively. The contacting sites are further delineated in Fig. 2E. **Fig. 2E:** A representation of the SARS-CoV-2 RBD structure. The critical contact sites of the amino acids are marked according to Wang *et al*. ^6^. **Fig. 2F:** The signal peptide sequence (SS) consists of 19 amino acids from the beginning. A cleavage site is at amino acid 13. A partial immunogenic epitope in the SS is marked in red letters. The glycan site, “NLT,” or “N” is marked with a blue highlight. Structure-based sequence alignments of the RBD related to **Fig. 2D and 2E**. The secondary structure elements are defined based on an ESPript ^21^ algorithm and labeled based on the RBD structure reported in the study. Dashed lines represent alpha strands and arrows represent beta strands. Three glycosylation sites (N) are highlighted in blue. Two antigenic epitopes, η1 and η2, are highlighted in yellow. Critical receptor binding domains consisting of essential amino acids are highlighted in green, and the number of positions of the amino acids is shown at the top.

### The signal peptide sequence (SS) positively impacts RBD secretion, location, and expression

Next, we studied spike protein and fragment expression by SDS–PAGE in an immunoblot assay using an anti-6x-his-tag mAb. SS-RBD and S1 exhibited higher expression than RBD and the full-length spike protein in the cell lysate at 24, 48, and 72 hrs p.t. (Fig. 2A). The RBD fragment and full-length spike protein were hardly detectable. In the cell pellet, RBD, SS-RBD, and S1 accumulated significantly; the full-length spike protein exhibited lower expression than the fragments; the S2 fragment was expressed in the cell pellet, but the yield of S2 was lowest among all expressed proteins in both cell lines (Fig. 3A, B). The vector control did not show any specificity in an immunoblot assay. We also observed that the full-length spike protein appeared in various sizes. This observation supported the concept that the spike proten constantly undergoes modifications, such as the addition of glycosylation chains or conformational changes, during its biological processing.

**Figure 3.**
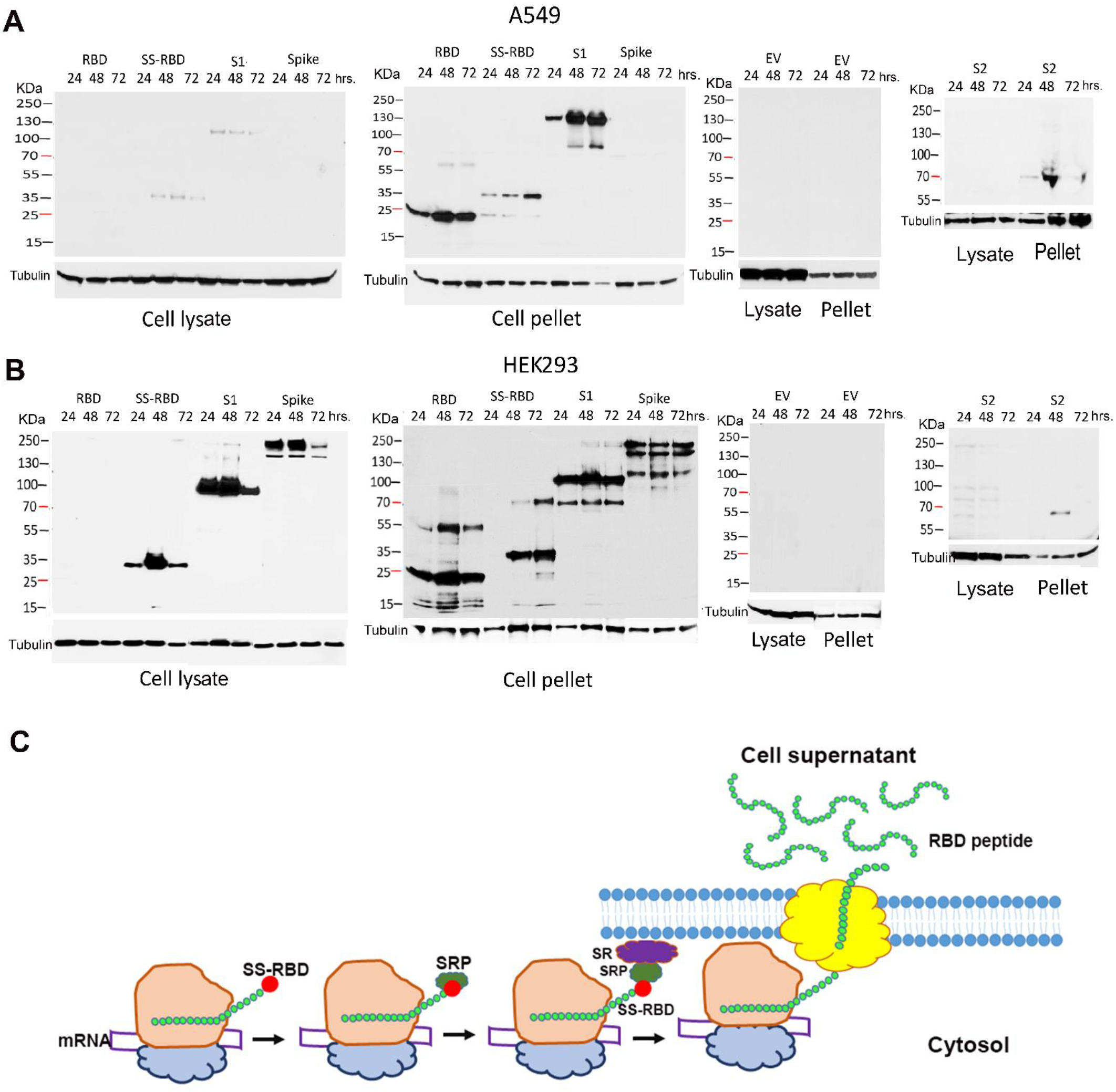
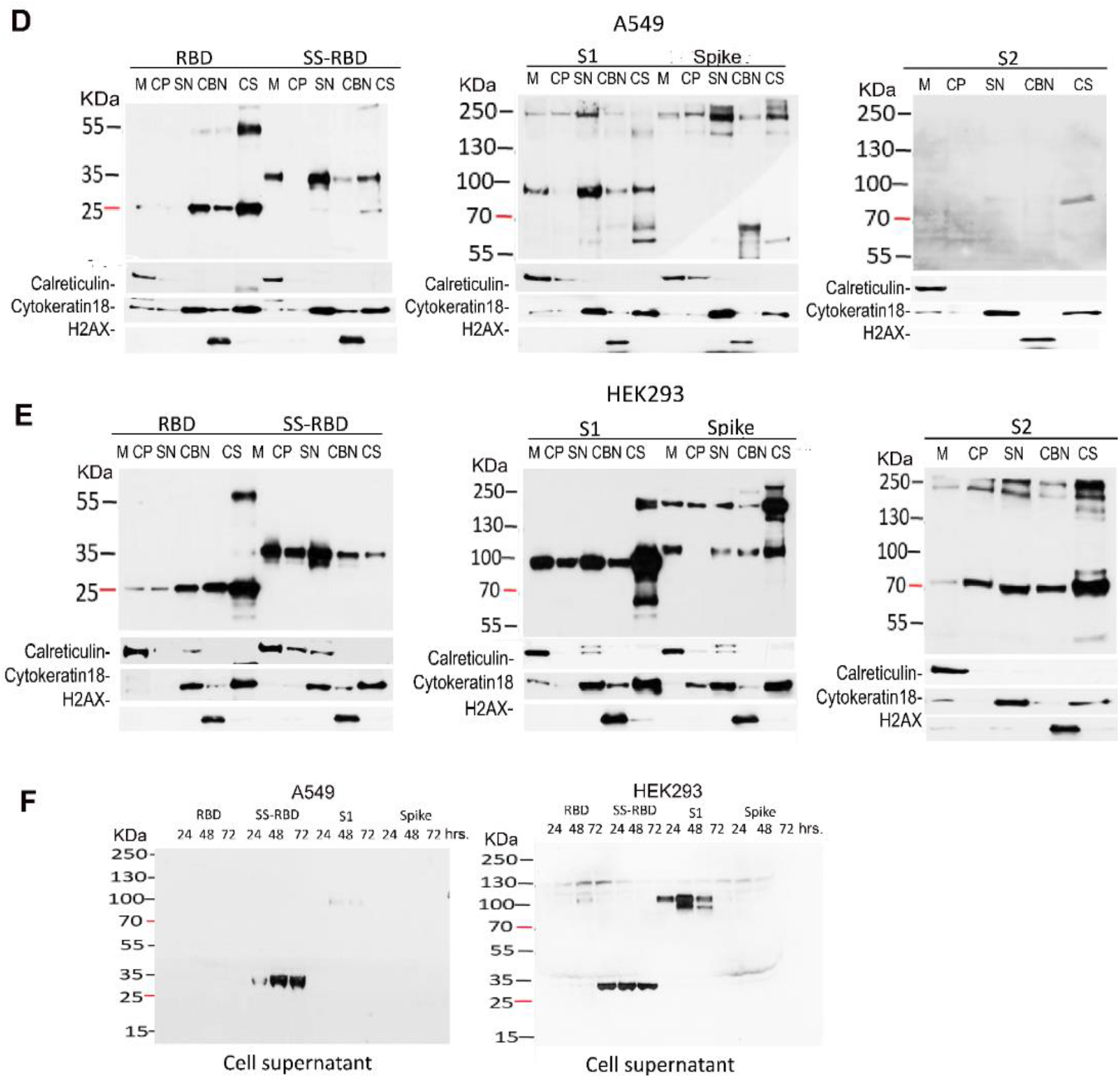
Full-length spike protein and fragments are present at different time points in cell lysates, pellet subcellular fractions, and cell supernatants in A549 and HEK293 cells. **Fig. 3A:** Kinetic expression of **full-length** spike protein and fragments in cell lysates, cell pellets, and subcellular fractions by immunoblotting. EV means empty vector. **Fig. 3B:** Western blot detection of spike protein fragments using an anti-His antibody at 24, 48, and 72 hr p.t. RBD (26 kDa), SS-RBD S1 subunit (100 kDa), and full-length spike (150 kDa), film exposure for 1 min. An anti-tubulin antibody detected tubulin as an internal reference in cell lysate and pellet samples. **Fig. 3C:** Illustration of the functional mechanisms of the signal peptide sequence in the SS-RBD, S1 subunit, and full-length spike in HEK293 cells. **Fig. 3D, 3E:** Immunoblot assay results of SARS-CoV-2 full-length spike protein and fragments in subcellular fractionated materials. A549 cells (Fig. 3 D) or HEK293 cells (Fig. 3 E) were fractionated by the Subcellular Protein Fractionation Kit. Normalized concentrations of each extract (10 μg) (the cytoskeleton extract concentration was 5 μg) were analyzed by Western blotting using specific antibodies against proteins from various cellular compartments, including calreticulin for detecting the membrane (M) and endoplasmic reticulum (SN) fraction, cytoskeleton marker (Cytoskeleton 18) for detecting the soluble nuclear (SN) fraction and cytoskeleton (CS) fraction, and rabbit anti-histone H2A. X mAb for detecting the chromatin-bound nuclear (CBN) fraction. A rabbit anti-His tag mAb detected the levels of the spike protein and fragments. Cell supernatants were harvested and denatured and then run on a mini-Nu-page gel. We extended the time to 2 hrs to visualize the protein band in A549 cells compared to 15 minutes in HEK293 cells. Thus, HEK293 cells exhibit higher SS-RBD and S1 protein expression levels than A549 cells.

### S1 and full-length spike protein were comparatively distributed in transfected cells

We used a thermal subcellular fractionation kit to elucidate the biological characteristics of the spike protein and fragments in transfected cells. In A549 and HEK293 cells, the RBD fragment was mainly expressed in the CS elution, followed by the SN and CBN elution (CBN). The distribution of SS-RBD was extensive compared to that of RBD. SS-RBD was mainly expressed in the CM, CP, and SNE elutions and was limited in the CBNE and CSE elutions. Therefore, immunoblot observed migration between the RBD and SS-RBD from the cytoskeleton to the cell surface (**Fig. 3B, 3C, 3D, 3E**). We also noticed that the spike protein and fragments were expressed higher in HEK293 cells than in A549 cells. S2 exhibited the lowest expression among all proteins studied; S2 peptides were identified in the CSE fraction in A549 cells. In contrast, S2 was distributed in all fractions except the CM fraction in HEK293 cells. In the CS fraction, spike protein and S1 subunits appeared as several bands, which represented biological processing and modifications. Only SS-RBD and S1 were detectable in the cell supernatant of the two cell lines at the three-time points (Fig. 3F).

### The PEA demonstrated that the spike protein and fragments were differentially expressed on a logarithmic scale

The PEA total expression, measured in the cell lysate and supernatant, was multiplied by the volume obtained for individual samples. We further analyzed the percentage of intracellular or extracellular distribution. The results showed that 70-80% of SS-RBD and S1 expression was released into the cell supernatant at 24 to 48 hrs p.t., and the expression rate increased to 94-97% at 72 hrs. p.t. In contrast, the full-length spike protein was expressed at 19%, 55%, and 70% in the cell supernatant at the three-timepoints. Moreover, a relatively high expression of the spike was observed in the cell lysate (Fig. 4A-E).

**Figure 4:**
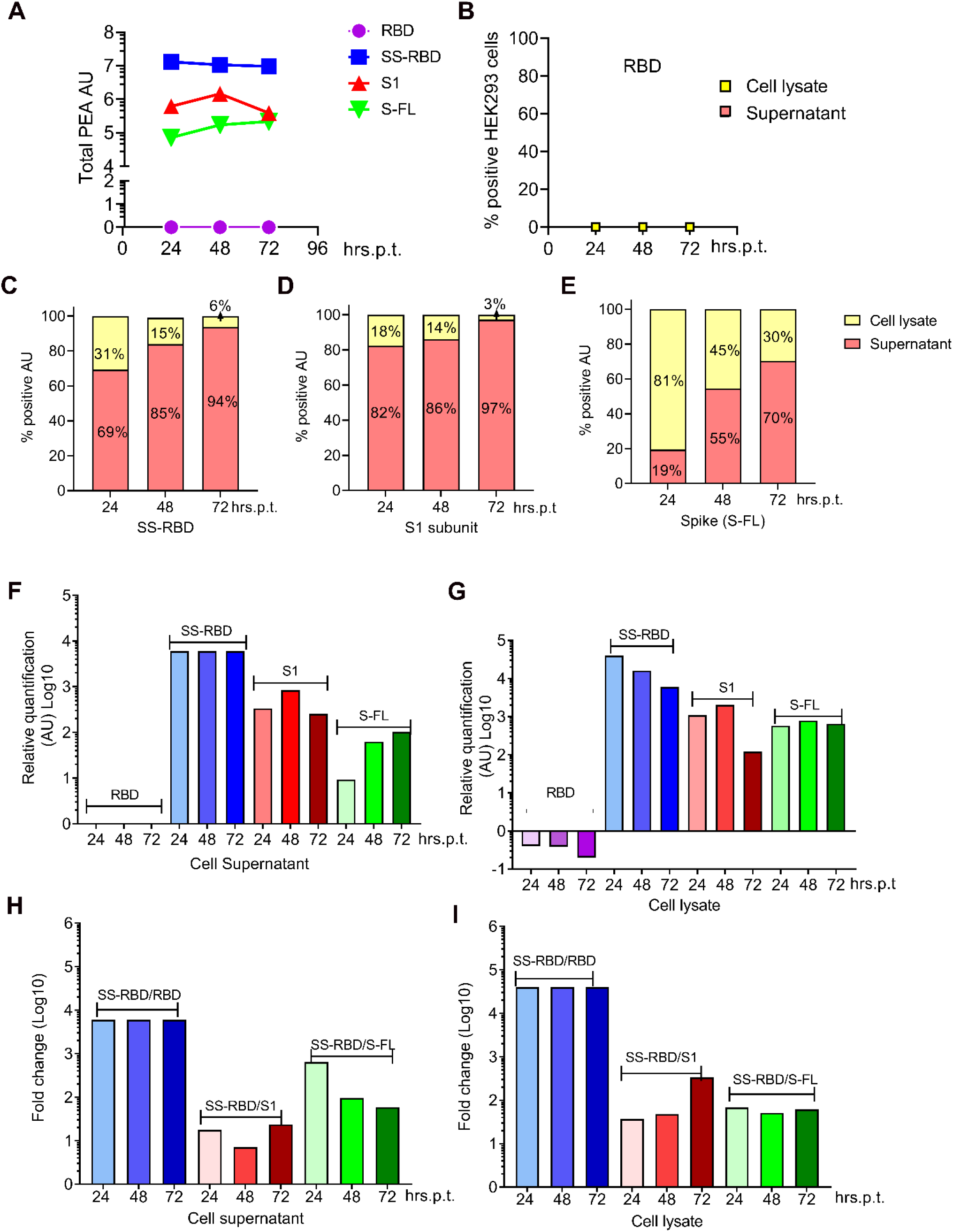
The expression levels in cell lysate and cell supernatants were quantified using highly sensitive proximity extension assays (PEA) in HEK293 cells at 24, 48, and 72 hrs p.t. Cell lysate and cell supernatant were serially diluted 10-fold and each dilution was duplicated. Finally, the results were considered reliable. A known quantity of RBD sample with a similar dilution was used as a quantity control. **Fig. 4A**: Total PEA relative quantification activation for spike proteins in cell supernatant and cell lysate from HEK293 cells. **Fig. 4B:** PEA relative quantification for the RBD domain. RBD exhibits an undetectable PEA activity unit (AU) due to being in the cell pellet. **Fig. 4C, 4D, 4E:** Total PEA relative quantification activation as AU for SS-RBD, S1 subunit, and the full-length spike protein, respectively. SS-RBD showed the highest expression (10^7^), the S1 subunit showed middle-level expression (10^6^), and the full-length spike protein showed middle-level expression (approximately 10^5^). Not shown: RBD was undetectable in the PEA because it was only expressed in the cell pellet. **Fig. 4F, 4G:** Comparison of the expression of the full-length spike protein and fragments in the cell supernatant and cell lysate, respectively. **Fig. 4H, 4I:** Fold change of SS-RBD expression over the expression of RBD, S1 subunit, and a full-length spike protein in cell supernatant and lysate.

The cell supernatant of HEK293 cells transfected with RBD was negative (Fig. 4B); the full-length spike protein was detectable at 100-fold dilution. The total S1 expression varied between 10^5^ and 10^6^ (AU) at 24, 48, and 72 hrs p.t. The expression of S1 was approximately one log higher than that of the full-length spike protein. The expression of SS-RBD was superior to all others and was expressed up to 10^7^ AU according to the PEA, which was approximately 100-fold more than the expression of the full-length spike protein in HEK293 cells (**Fig. 4C, D, E**).

The PEA demonstrated that SS-RBD expression is superior to the S1 fragment and the full-length spike protein in the HEK293 supernatant. RBD expression was lower than the standard curve range at the three-time points, so it was undetectable, consistent with the above FACS data. Notably, two different Nabs negatively detected RBD antigens in the PEA and FACS experiments. However, RBD expression was negative in both assays. Our results supported that ACE2 completely blocked RBD antigenicity.

### Confocal microscopy demonstrated the expression of S1 and the full-length spike protein in both the cytoplasm and nucleus

A confocal microscope was used to monitor the expression and localization of the spike protein and fragments in A549 and HEK293 cells. At 24 hrs p.t, A549 and HEK293 cells were subjected to immunofluorescence staining; the plasmids containing the RBD, SS-RBD, and S2 fragments were expressed in the cytoplasm, whereas the plasmids containing the S1 fragment and full-length spike protein were detectable in both the cytoplasm and cell nucleus. No transfected cells or empty vector-transfected cells did not show any fluorescence. The biological distributions of S1 and full-length spike protein were consistent in A549 and HEK293 cells (**Fig. 5A, 5B**). The percentage of positively stained cells was determined under standard fluorescence microscopy. The positively stained cells were counted at least four times in different areas. In comparing the five DNA plasmids, SS-RBD exhibited the highest expression among the five spike proteins, S2 expression appeared lowest, and RBD, S1, and full-length spike protein were expressed at mid-range levels (**Fig. 5C, 5D**).

**Figure 5.**
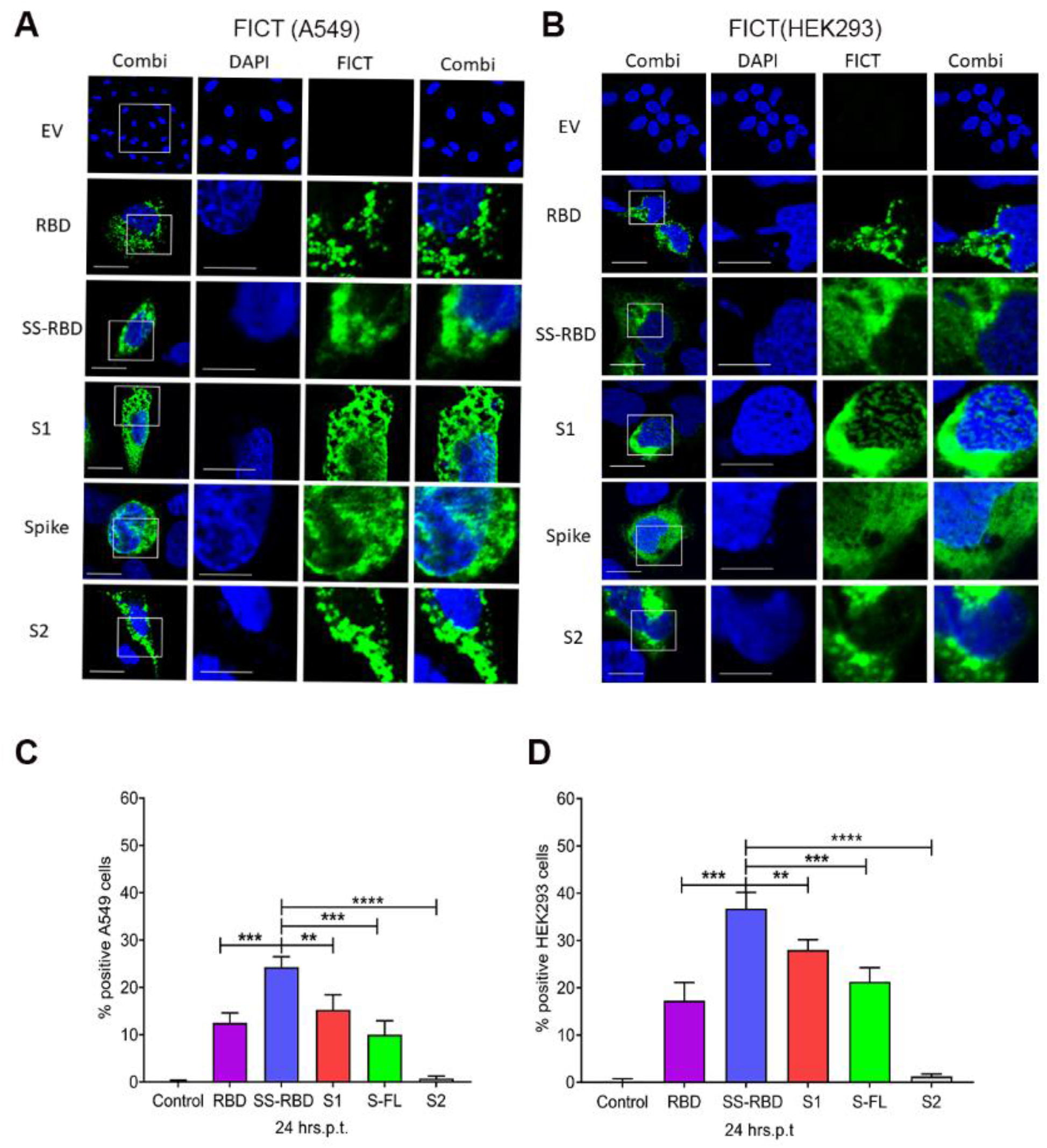
Confocal microscopy imaging for the localization and expression of the full-length spike protein and fragments at 24 hrs p.t. in **Fig. 5A, 5B**. A549 and HEK293 cells were grown on glass coverslips overnight; the cells were transfected with plasmid DNAs, incubated for 24 hrs, fixed, and processed for immunofluorescence staining (materials and methods). The full-length spike protein and fragments’ subcellular distribution was stained with an anti-6-his tag mAb and FICT secondary Ab (green), and nuclei were stained with DAPI (blue). Bar: 10 μm. **Fig 5C, 5D**: A549 and HEK293 cells were also calculated as the percentage of cells positively transfected for the full-length spike protein and fragments under immunofluorescence microscopy. The graph presents the averages from triplicate samples. The asterisks indicate significant differences, as demonstrated by an unpaired T-test (* P < 0.05; **P < 0.01; ***P < 0.001); ns, not significant. The data are presented as the mean ± SE from three samples.

## Discussions

mRNA-based SARS-CoV-2 vaccines have been globally dispersed to prevent infection and disease. The RBD of a spike protein is an imperative target for serological and B-cell studies because it directly binds ACE2, mediating host cell entry ^13 22^. Usually, antibodies interacting with the RBD potently block the binding of the virus to ACE2 and thereby neutralize the virus (Barnes *et al*., 2020). One of the candidate strategies for vaccine development is to develop a protective vaccine through the induction of antibodies against the RBD domain ^23^. In the present study, ACE2 could bind to the RBD on the spike protein, which blocked the RBD from polyclonal NAb detection. Our findings explained why most of the isolated mAbs did not recognize the RBM, and the mAbs that neutralized authentic SARS-CoV-2 were unable to inhibit spike protein binding to ACE2. mAbs against RBD might inefficiently neutralize or block the viral interaction. Atomic ultrastructure analysis demonstrated that the epitopes recognized by Nab are colocalized with the interface between RBD and ACE2, blocking antigenicity. Fortunately, SS-RBD contains an additional glycan at N17 in the SS peptide that partially unblocks the antigenicity of RBD, resulting in NAb recognition ^24^. Although non-neutralizing antibodies induced by mRNA vaccines can reduce viral infection at a certain level, the RBD on the viral surface still binds to ACE2, which leads to viral internalization into host cells. Consequently, the virus can cause reinfection. Our study suggests that reinfection can occur in mRNA-vaccinated populations because of the blocked antigenicity of the RBD.

Early experiments using RBD in mRNA spike protein vaccines induced accessory and efficient NAbs via ACE2 affinity. mRNA vaccination appeared to cause a high antibody response of relatively homologous viral titers. However, a large study showed that vaccines generate more non-neutralizing antibodies than COVID-19 survivors do, resulting in a lower ratio of neutralizing to binding antibodies ^25^. Several reports have demonstrated that non-neutralizing antibodies also have a protective effect against many viral infections, especially when neutralizing antibody titers are very low or absent. In addition, natural immunity induced by the mRNA vaccine is substantial and protects against virus infection ^10^. Notably, a clinical report showed that 83,2% of patients suffering from severe SARS-CoV-2 also exhibited lymphopenia ^26^. It is still unclear whether a spike protein interaction might cause lymphopenia. Recent studies have demonstrated that the spike protein can cause severe adverse effects in young adults, disrupt cardiac pericyte function, and trigger multiple necrotic encephalitis ^11,27^. We showed that the blocked RBD antigenicity on the spike protein suppresses the production of NAbs against RBD. Therefore, we speculated that as a candidate for a new mRNA vaccine instead of the current only RBD, that is not a suitable, potent, or stable immunogen for in vivo human trials.

The immunogenicity of the studied spike protein fragments is distinctly different between in vivo animal models and humans. Several reports have shown that vaccines targeting the RBD induce protective immunity. Mice, rabbits, and nonhuman primates (*Macaca mulatta*) immunized with various sizes of the spike protein can produce two- to fivefold-fold more RBD antibodies than those of other spike variants. Furthermore, vaccination in nonhuman primates shows that an in vivo challenge with SARS-CoV-2 viruses can reduce the viral load in the lung of mice model^28 29^. We observed that the spike protein fragments acted as potent immunogens in animal model experiments. However, they served as both receptor-binding proteins and immunogens in humans. Thus, we speculated that the duality of the spike protein in humans may be the main challenge for the induction of non-neutralizing antibodies. Consequently, the protective effect of the spike mRNA vaccine on model animals cannot fully support its effectiveness in humans, owing to the spike’s duality.

Unlike classical protein vaccines that act as immunogens and directly reach their highest concentration for stimulating immune cells and inducing antibodies, spike mRNA/DNA vaccines must use their genetic materials to enter host cells and time-dependently express and gradually release immunogens from target cells. Moreover, the newly generated spike serves as not only an immunogen but also binds to ACE2 in target or neighboring cells. Besides the ACE2 interaction, we cannot rule out the possibility that hydrophobic interactions could also block RBD antigenicity, although the hydrophobicity is not strong. To our knowledge, virus binding occurs much quicker than the immune system can recognize. Therefore, the blocked RBD can, but the exposed surrounding RBD cannot, escape the recognition of macrophages and B cells producing NAbs.

The RBD comprises 220 amino acids (aa), and the RBM (74 aa) is 1/3 of the RBD. 8-10 aa in the RBM plays critical roles in ACE2 binding affinity and potent immunogenicity. The exchange of one or a few amino acids in the RBM of new variants can reduce the affinity for ACE2 and increase the recognition by NAbs ^25^. Removing ACE2 affinity may reveal RBD antigenicity. Moreover, it could be the basis of a universal vaccine against SARS-CoV-2 variants since the most dominant variations occur in the RBM region ^15^. Although abolishing ACE2 binding affinity may not result in increased recognition by NAbs to prevent reinfections, it could, at the very least, reduce the adverse effects of RBD-ACE2 interaction in vivo ^11^. In the context of vaccine development, abolishing ACE2 binding affinity on the spike protein might also lose some immunogenicity of the RBM but could gain the majority of immunogenicity, approximately 70% of the RBD. In SARS-CoV-2 variants, the reduced affinity of E484K for hACE2 demonstrated that it is more susceptible to RBD-binding mAbs. These findings support our proposal of modifying the interaction between ACE2 and the spike protein to gain more stable, potent, and effective antigenicity than the presently used spike protein vaccine. Moreover, the modification could be beneficial for avoiding spike protein-induced DNA damage to endothelium proteins and impaired mitochondrial function caused by ACE2 downregulation ^30^.

In the present study, we studied the biological characteristics of six variants of SARS-CoV-2 in A549 and HEK293 cells using four different technologies. We demonstrated that the RBD fragment is efficiently expressed and mainly located in the cytoplasm, bound together with the cytoskeleton and that SS-RBD, S1, and the full-length spike protein were detected in the cell supernatant at different levels (Table 1). Western blot analysis demonstrated that SS-RBD manifests in the majority of cell lysates and cell membranes (Fig 4). Kinetics of the spike protein and fragments showed that SS-RBD and S1 displayed high expression in the intracellular phase and cell supernatant; the full-length spike protein exhibited upregulated expression on the cell surface; the S1 subunit and full-length spike protein demonstrated similar localization throughout the cell, such as in the cell lysate and cell pellet as well as five subcellular fractions. The S2 fragment, which makes up more than half the C-terminus of the spike protein, containing a transmembrane domain, was expressed at a low level in transfected kidney and lung cells. S2 was tightly bound to the cytoskeleton and located in the cytoplasm.

**Table 1:**
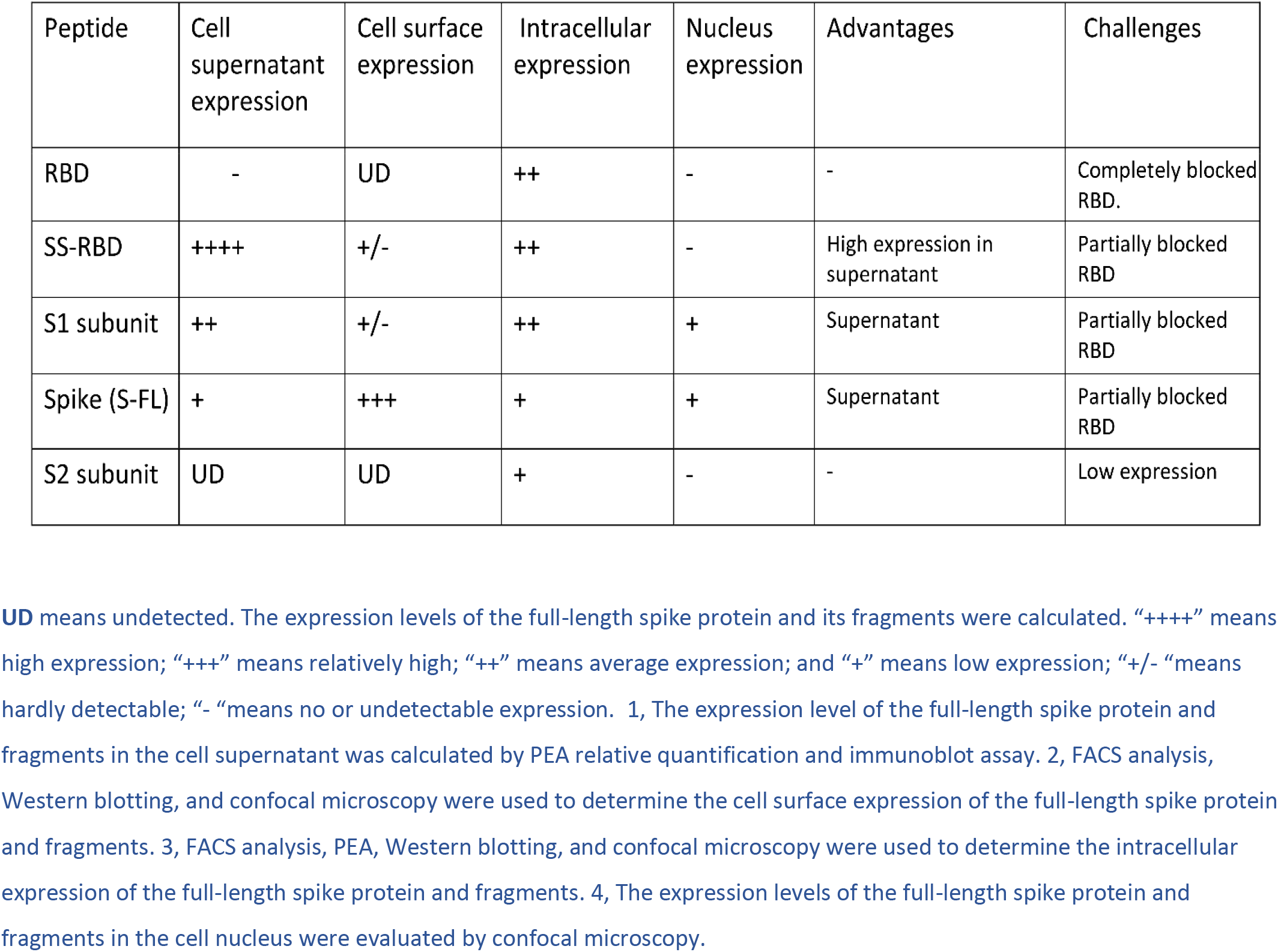
Secretion, expression, and location of the SARS-CoV-2 full-length spike protein and its fragments revealed advantages and challenges for mRNA vaccine efficiency

The PEA method consists of three main steps: a liquid antigen brings oligonucleotide-conjugated antibody molecules into proximity, extension/pre-PCR, and readout via real-time PCR. The reaction used 1 μl of undiluted or 10-fold diluted sample. The antigen was measured up to 7 logs for SS-RBD, 6 logs for S1, and 5 logs for the full-length spike protein. Thus, PEA is sensitive, stable, and has a broad dynamic range for RBD antigen detection. We demonstrated that the full-length spike protein was undetectable in immunoblot but could be detected in PEA after 100-fold dilutions, demonstrating that PEA was superior to immunoblot for antigen detection. PEA quantification showed a 100- or 10-fold increase in SS-RBD or S1 levels compared to full-length spike protein levels (Fig. 4). However, RBD without the SS was not detected with the NAb in either the immunoblot or PEA method. The SS stimulated the movement of its peptide from the cytoskeleton toward the cell surface until it was released into the cell supernatant.

Notably, the first 13 amino acids can function sufficiently as the SS, as reported by Liu *et al*.^31^; in the present study, the SS consisted of 19 aa with an additional glycan at N17 covering a partial antigenic epitope that can be recognized by NAb ^24^. SS-RBD can be identified by the signal recognition particle (SRP), which is a ribonucleoprotein complex comprising six polypeptides (>100 aa/each) and one SRP RNA. After the SS-SRP complex moves along the ER, the SRP binds to the SRP receptor (SR), and then SS-RBD is released into the cell supernatant. The SRP has an enormous molecular weight, as it interacts with the hydrophobic SS that affects RBD conformation, revealing antigenicity in SS-RBD. Thus, the SS we designed improved truncation and antigenicity. In contrast, the RBD and S2 fragment lack SS. They could not migrate to the membrane destination and were retained in the cell cytoplasm, more precisely in the cytoskeleton, and ER markers were detectable (Fig. 3D. E). Other reports also support the notion that mAbs outside of the RBD, such as S309 and N343, have a neutralizing function^32,33^.

A protein’s nuclear localization signal (NLS) is required to translocate the cytoplasm and nucleus. More than a dozen NLSs with various scores throughout the spike protein were identified by a computer program. Moreover, most motifs are localized in the S1 domain; one or more predicted NLSs might trigger the S1 fragment or spike protein to enter the nucleus. Ultrastructure analysis in an early SARS-CoV-2 infection model showed that the annulate lamellae (AL) were close to the cell nucleus ^34^. AL is newly formed only in cells with active viral replication, and these cytoplasmic structures present a site of viral glycoprotein accumulation ^35^. The nuclear pore compartments (NPCs) of AL could potentially mediate exchanges between these partially sealed compartments and the cytoplasm, as demonstrated in herpes virus and hepatitis C virus infection ^36^. In this study, S1 was highly aggregated around and in the nucleus under confocal microscopy (unpublished data). Therefore, we assumed that the potential mechanisms of S1 in the nucleus might be via NPCs, which was recently clarified by Sattar *et al*. ^37^.

A promising vaccine should be stable, effective, and practical. A low dose should induce high efficiency NAbs that maintain a long protective period, such as hepatitis A and B and polio vaccines. If a spike protein had a partially abolished RBM, the remaining immunogenicity might not sufficiently stimulate NAbs to fight against the viral invasion of a new variant. Moreover, not all pathogens can make effective or applicable vaccines. As a result of blocked RBD antigenicity, only the surrounding RBDs on the spike protein are recognized by immune cells, inducing weak and unstable antibodies that quickly degrade. In the early days of HIV research, people were optimistic that a single gp120 molecule would be efficient enough to make a vaccine. Nevertheless, the development of an HIV vaccine just recently had a breakthrough. For an effective SARS-CoV-2 vaccine, more attention to basic virology and immunology is necessary and imperative before producing a large-scale vaccine.

In conclusion, our findings indicate that NAbs can be used to detect SS-RBD but not RBD. ACE2 receptors block antigens of RBD in transfected cells, so using RBD or SS-RBD as immunogens for mRNA vaccines may be insufficient to protect against SARS-CoV-2 infection due to unexposed antigens on the RBM. Therefore, future vaccines need to eliminate ACE2 binding affinity and expose antigenicity on the RBD. Furthermore, identifying critical amino acids involved in nuclear localization and reducing nuclear entry could improve spike protein expression, translation, and secretion and reduce underlying pathogenesis in the nucleus. In the past, reports had claimed that the neutralizing effect of some potent mAbs was reduced or absent when a new variant appeared. Thus, scientists have long struggled to detect unknown variants as early as possible to prepare an updated vaccine. Our results elucidate that complete blocking of the RBD is a pivotal contributor to the dysfunction of potent NAbs. Therefore, a spike protein lacking ACE2 affinity is required for new mRNA vaccines. Thus, our findings provide the field with new insights into the biological and immunological functions of the SARS-CoV-2 spike protein and the development of new vaccines against the spike protein.

## Declaration Acknowledgments

In this article, Fig 2C and 2D were adapted from an imperative article published by Qihui Wang, the first author and the leading author Jianxun Qi and the first author of “ Structural and functional basis of SARS-CoV-2 entry by using human ACE2” in Cell, 2020 (https://doi.org/10.1016/j.cell.2020.03.045)”. We acknowledge the Cell Press *Elsevier” and the authors for the permission to use the illustrations. We want also to thank the Biochemical Imaging Center at Umeå University and the National Microscopy Infrastructure, NMI (VR-RFI 2019-00217) for assisting in microscopy.

This research was funded by the Umeå University Medical Faculty’s Planning grants for COVID-19 (research project number: 3453 16032) awarded to Dr. YF Mei and ALF-Basenhet in 2020 and 2021 and Dr. YF Mei in 2022, and the Lion’s Cancer Research Foundation at Umeå University (grants AMP 19-982, LP20-2256, and AMP22-1071 awarded to Dr. YF Mei).

## Author Contributions Statement

QZ, ZWF, and YFM conducted the plasmid preparations, FACS assays, and immunoblotting. SJ, QZ, and YFM conducted the confocal assay. MH and HZ conducted the PEA experiments and reviewed the manuscript. YFM planned the experiments and wrote and reviewed the manuscript. YFM and ML supervised the study.

## Notes

### Competing Interest Statement

The authors have declared no competing interest.

